# NOCICEPTRA2.0 - a comprehensive ncRNA atlas of human native and iPSC-derived sensory neurons

**DOI:** 10.1101/2023.04.24.536852

**Authors:** Maximilian Zeidler, Diana Tavares-Ferreira, Jackson Brougher, Theodore J. Price, Michaela Kress

## Abstract

Non-coding RNAs (ncRNAs) play a critical role in regulating gene expression during development and in the pathogenesis of diseases. In particular, microRNAs have been extensively studied in the context of neurogenesis, the differentiation of pain sensing nociceptive neurons and the pathogenesis of pain disorder, however, little is known about the developmental signatures of other ncRNA species throughout sensory neuron differentiation. Moreover, there is currently no information available about the general expression signatures of ncRNAs in human dorsal root ganglia (DRGs) harboring the cell bodies of primary afferent nociceptors.

To bridge this knowledge gap, we developed a comprehensive atlas of small ncRNA species signatures during the differentiation of human induced pluripotent stem cell (iPSC)-derived nociceptive neurons. By employing a combination of iPSC-derived sensory neuron and human DRG long and short RNA co-sequencing, we identified specific signatures that describe the developmental processes and the signatures of all currently known small ncRNA species in detail.

Our analysis revealed that different ncRNA species, including tRNAs, snoRNAs, lncRNAs, and piRNAs, are associated with different stages of sensory neuron differentiation and maturation. We retrieved pronounced similarities in ncRNA expression between human DRG and late-stage iPSC-derived sensory neurons, which further supports the use of iPSC-derived sensory neurons to uncover functional and regulatory changes in ncRNAs and their suitability as a as a human model system to bridge the translational gap between preclinical findings mostly from rodent models and our understanding of human disorders for the development of mechanism-based treatments.

In summary, our findings provide important insights into the role of ncRNA species other than microRNAs in human nociceptors. The updated NOCICEPTRA2.0 Tool will be the first fully comprehensive searchable ncRNA database for human sensory neurons enabling researchers to investigate important hub ncRNA regulators in nociceptors in full detail.

## Introduction

Small non-coding RNA (ncRNA) species are increasingly emerging as important regulatory hubs in health-related biological processes and the pathogenesis of a multitude of diseases including pain disorders. These short ncRNA species include microRNAs (miRNAs), small nucleolar RNAs (snoRNAs), small nuclear RNAs (snRNAs), transfer RNAs (tRNAs) and ribosomal RNAs (rRNAs), PIWI-interacting RNAs (piRNAs) and small interfering RNAs (siRNAs), ranging from smaller molecules such as mature miRNAs (∼22 nucleotides) to full length sno- and tRNAs (∼70-120 nucleotides) (Aalto & Pasquinelli 2012). While initially neglected as “*junk RNA*” fragments with no functionality, relevant functional roles of the different ncRNA classes are increasingly uncovered, such as the degradation of transcribed messenger RNA products via direct interaction, and the regulation of protein translation or mRNA splicing through their interaction with ribosomal proteins or spliceosomes (Aalto & Pasquinelli 2012; Iwasaki *et al*. 2015; Kufel & Grzechnik 2018). Species and tissue specific expression databases based on high-throughput next-generation sequencing show that ncRNAs are specifically distributed and expressed in cell types and tissues suggesting that ncRNAs drive specific cell type functionality (Godoy *et al*. 2018; Isakova *et al*. 2020; Stefani & Slack 2008). In addition, fragmented mature ncRNAs such as tRNAs and snoRNAs can be incorporated into the Argonaute complex thereby mimicking the function of miRNAs (Guan *et al*. 2020; Kumar *et al*. 2014). For example, upregulation of tRNA fragments inducing neuronal necrosis is detectable in post stroke patient blood samples (Cao *et al*. 2021; Winek *et al*. 2020).

Dramatic changes in the expression of coding genes, ncRNAs, pseudogenes, and splice isoforms are seen during the transition from pluripotent stem cells to early differentiating neurons and several lncRNAs undergo dramatic expression changes suggesting important roles for ncRNAs in neurogenesis (Hjelm *et al*. 2013; Lin *et al*. 2011). Distinct ncRNA expression signatures are retrieved from different cell types indicating that ncRNA signatures can represent and resolve a particular phenotype (Godoy *et al*. 2018; Isakova *et al*. 2020; Zhao *et al*. 2016). However, cell specific ncRNA signatures are still in their infancy and temporal developmental dependencies of cell-type specific markers derived from single cell and bulk RNA sequencing studies are under-investigated or even missing for ncRNA species other than miRNAs. ncRNA transcriptome analyses are usually more complex compared to mRNA sequencing approaches due to relevant caveats in the alignment and counting process of ncRNAs, mostly correlated with shorter nucleotide length and the tendency of multiple occurrences and annotations within the genome. This specifically applies to shorter variant ncRNA species such as tRFs, snoRNAs, piRNAs and miRNAs (Deschamps-Francoeur *et al*. 2020). Furthermore, alignment rates of small RNAs in scRNA studies are poor because of the high adapter content and increased degradation of short RNA variants (Faridani *et al*. 2016; Hagemann-Jensen *et al*. 2018; Hücker *et al*. 2021). Thus, our understanding of ncRNA signatures and their roles throughout human neuronal development remains limited, and neither temporal single cell transcriptomic data nor bulk-ncRNA transcriptomic datasets specific for human neuronal development are available.

Nociceptors are primary afferent sensory neurons serving the transduction of noxious stimuli leading to the perception of pain. Although rodent models are currently the *gold-standard* for functional sensory neuron research, single cell RNA studies identified substantial differences between human and rodent sensory neurons, indicating sex- and species-specific functionalities with potential relevance for the forward and reverse translation of findings from rodent models towards human pain disorders (Tavares-Ferreira *et al*. 2022; Zeidler *et al*. 2022). Recent advances in the reprogramming of induced pluripotent stem cells (iPSC) allow to develop human iPSC derived nociceptors which offer improved opportunities for modelling human pathophysiologies in depth (Chambers *et al*. 2012; Schoepf *et al*. 2020).

Developmental trajectories of mRNAs and microRNAs (miRNAs) characterize five different stages of nociceptor differentiation from human iPSC (Zeidler *et al*. 2021) and likely play important roles in developmental processes via direct or indirect regulatory pathways. Tissue and cell specific expression of ncRNAs is associated with cell type functionalities, phenotypes and morphologies, however, mechanistic insight into their functionality remain sparse (Isakova *et al*. 2020). Despite their critical importance for health and disease, developmental signatures of ncRNAs are mostly explored for miRNAs (Lundin *et al*. 2020; Zeidler *et al*. 2021) whereas unbiased knowledge on other ncRNA species relevant for human nociceptor development is currently not available. Therefore, we for the first time provide a comprehensive atlas of ncRNA signatures, that potentially contribute to human iPSC-derived nociceptor development and maturation, with a specific focus on individual ncRNA expression trajectories, their importance in specific developmental stages and ncRNA::mRNA connectivity.

## Methods

### Data Fetching and iPSC-derived sensory neuron differentiation

Long RNA and small RNA datasets were retrieved from GEO Datasets with the Accession ID: GSE161530. Details of the experimental sequencing procedures as well as the induction of induced pluripotent stem cell (iPSC) differentiation into nociceptive neuron like cells are described in (Schoepf *et al*. 2020; Zeidler *et al*. 2021). Briefly, iPSCs were seeded as single cells on Matrigel coated 6-wells (8×10⁵ cells) and maintained in mTeSR medium supplement with Rock Inhibitor to increase cell survival. AD2, AD3, and 840 iPSCs were cultured on Matrigel (Corning) coated 6-well plates and maintained in mTeSR1 medium (STEMCELL Technologies). Nociceptor differentiation was performed as previously published.

In brief, ≈80× 10^4^cells per 6-well plate were seeded as single-cells at passage 2 after thawing. Cells were maintained overnight in mTeSR1 medium containing the Rho-associated kinase (ROCK) inhibitor Y-27632 (Sigma Aldrich) until cells reached 60–70% of confluency. Subsequently, dual-SMAD inhibition was initiated by inhibiting BMP4 and TGF-β in knockout serum replacement (KSR; Knockout DMEM [Gibco], 15% KO serum replacement [Gibco], 1% non-essential amino acids (PAA), 2 mm Glutamax [Gibco], 1× Penicillin/Streptomycin [Sigma Aldrich], and 100 µm β-mercaptoethanol [Gibco] medium supplemented with 100 nm of LDN-1931189 (STEMGENT) and 10 µm SB431542 (selleckchem.com) for 5 days. Additionally, 3 µm CHIR99021 (GSK3beta inhibitor, selleckchem.com), 10 µm DAPT (gamma-secretase inhibitor, selleckchem.com) and 10 µm SU5402 (VEGFR-2, FGFR-1, and PDGFRB inhibitor, selleckchem.com) were added from day 2 (D2) to D12. From D4 on N2/B27 (Neurobasal medium/GLUTAMAX/2% B27 w/o vitamin A [Gibco]/1% N2 [Gibco] was added to the KSR medium every second day in increments of 25%. On D12, 80 × 10^4^ cells were passaged onto Matrigel coated 6-well plates according to the supplier’s recommendations and maintained in N2/B27 medium supplemented with 25 ng mL^−1^ nerve growth factor (hNGF, PreproTech) 25 ng mL^−1^ brain derived neurotrophic factor (hBDNF, PreproTech) and 10 ng mL^−1^ glia derived neurotrophic factor (hGDNF, PreproTech). Y-27632 was added for 1 day following passaging, to promote survival of neuronal-like progenitor cells. Cytosine-β-D-arabinofuranoside (4 µm, Sigma Aldrich) was administered on D14 for ≈24 h to reduce the amount of non-differentiated proliferating cells. Cultures were maintained until D36 and medium was changed every other day.

### Human native DRG analysis

Recovery of human DRG samples derived from organ donors (**supplementary Table S1**) was done through a collaboration with Southwest Transplant Alliance and all protocols were approved by the Institutional Review Board of University of Texas at Dallas. RNA was isolated from lumbar and thoracal human DRG samples and total RNA was sequenced using the Illumina TrueSeq sequencing kit. Fastq files were aligned against the human transcriptome provided by Gencode using Salmon. Obtained transcript per million reads were transformed into gene aggregated row counts using the tximport module. Libraries were normalized using DESeq2 and variance stabilized counts retrieved for lncRNAs and correlated against all the individual timepoints of iPSC-derived sensory neuron development, only considering lncRNAs.

In addition, ncRNA libraries of human organ donors DRG samples were generated using the Nextflex Small RNA-Seq kit (supplementary Table S2). In short ncRNA fastq files were quality controlled using FastQC and trimming of adapter as well as random 4-bp long nucleotides trimming from each end of the read were performed using cutadapt with parameters suggested by the Nextflex kit. Processed fastq files were then aligned and counted using the exceRpt pipeline against the hg38 reference genome and DESEq2 was used for count and library size normalization as well as generation of variance stabilized counts (vst). Finally, vst and normalized counts were compared with counts obtained from the iDN analysis.

### RNA read processing and differential gene expression analysis

Derived mRNA FastQ files were pseudo aligned against the human transcriptome (Genecode.v39.transcripts.fa) provided by GENCODE (www.gencodegenes.org) using *Salmon* (v0.13.1) software with the following parameters (-l A, -p 16, --numBootstraps 30, --useVBOpt –seqBias --validateMappings). Small RNA sequencing libraries were prepared using the NEBNext Small RNA Library Prep Set for Illumina as previously described (Zeidler *et al*. 2021). Small RNA reads were further processed and adapter sequences trimmed off (AGATCGGAAGAGCACACGTCTGAACTCCAGTCAC) using *Flexbar* v3 and subsequently aligned against the human reference genome using the exceRpt pipeline and the Manatee Pipeline(Dodt *et al*. 2012; Handzlik *et al*. 2020; Rozowsky *et al*. 2019). Reads were aligned and counted using the exceRpt pipeline using the default parameter order aligning first to miRNAs, tRNAs, piRNAs and then against the genome and circRNAs.

### Differential Gene Expression Analysis mRNA

Salmon retrieved mRNA quant files were aggregated on gene level counts using the tximport (v1.20.09) package and the Genecode.v39.annotation file. Differential gene expression was performed using the model ∼timepoint + cell line + timepoint:cell line, where time represents the different timepoints throughout development, cell line one of the three iPSC clones (AD3, AD2 and 840) and the interaction and compared this using the likelihood ratio test against a reduced model of ∼cell line. Log2Fold changes acquired by DESeq2 were shrinked using the “ashr” function and genes were considered differentially expressed following FDR-correction if the q-value is below 0.05. Quality Control of dispersion and principle component analysis (PCA) were performed based on variance stabilized counts, and lncRNAs were retrieved using the gene biotype annotation from the Gencode.v39.annotation file (Love *et al*. 2014).

### Differential ncRNA expressions

Count table with all expressed ncRNAs was used for differential ncRNA expression analysis in DESseq2 using the same model as described in the mRNA part. All ncRNAs were used for DESeq2 normalization and split into the individual ncRNA biotypes after differential gene expression, to access the time dependency using principal component analysis for each biotype individually. In addition, tRNAs were further aligned and counted using MINTmap which explicitly separated the specific tRNA fragment (tRF) types (5’, 3’, i-TRf etc.), which allows us to investigate tRF distribution in depth (Loher *et al*. 2017).

### Clustering of ncRNA trajectories

Variance stabilized counts of the differentially expressed ncRNAs for each biotype were used for hierarchical clustering analysis using the ward algorithm and dissimilarity (1-correlation) as distance metric for each ncRNA biotype. Individual trajectories were then summarized to a main trajectory of the module and visualized using a boxplot/lineplot combination implemented in pythons v3.9 “seaborn” package. Furthermore, ncRNAs were ranked based on the negative log10 FDR-corrected p-value, and top three dysregulated ncRNAs per cluster were determined and visualized.

### Databases

To depict functional changes of ncRNAs relevant databases were queried. For snoRNAs the snoDB (http://scottgroup.med.usherbrooke.ca/snoDB/) was queried and Box type (H/ACA or C/D) as well as the host-gene were joined and mapped to the respective differentially expressed snoRNAs (Bouchard-Bourelle *et al*. 2019). SnoRNAs were grouped by the hierarchical cluster and aggregated based on the percentage of Box types per hierarchical cluster. Host-gene expression for the differentially expressed snoRNAs were retrieved from our mRNA dataset and correlated against the expression of the individual snoRNAs to determine putative co-expression networks. Enrichment analysis of all positively and negatively correlated (corr. Coeff.> abs(0.5)) were used for g:Profiler enrichment analysis in the annotation spaces (GO:BP, GO:MF, GO:CC and KEGG) (Raudvere *et al*. 2019). LncRNAs and lincRNAs were joined manually based on the same cluster expression patterns, and a curated list was used to query miRNA target enrichment per hierarchical cluster using lncSEA (http://bio.liclab.net/LncSEA/index.php, (Chen *et al*. 2020)). Only significantly enriched miRNAs were used for further analysis and the top 5 enriched miRNAs per timepoint visualized by means of the negative log10 adjusted FDR-corrected p-value. Trajectories were additionally also retrieved from the manatee counts and transformed to compare.

### WGCNA analysis

WGCNA is a method used to identify groups of highly correlated genes or non-coding RNAs (ncRNAs) and to investigate the relationships among these groups in a biological context using their color-coded module assignments.All ncRNAs with a DESeq2 determined Base Mean of >10 were subject to the weighted gene correlation network analysis (WGCNA). VST normalized counts were subjected, and integrity of the data was determined using the goodSamplesGenes function integrated into the WGCNA package (Langfelder & Horvath 2008). Since WGCNA analysis should full-fill scale-free topology, the softThreshold was determined to be above 0.8. Data was then subjected to the blockwiseModule function to construct a signed network using the following parameters (power = 14, maxBlockSize = 20000, TOMType = “signed”, minModuleSize = 30, reassignThreshold = 0, mergeCutHeight = 25, numericLabels = False, pamRespectsDendro = False, saveToms = True, verbose = 3, randomSeed = 1234, networkType = “signed”, deepSplit = 2). Module affiliation for each gene was determined using the kME (k-Module Eigenvector affiliation). Hub-genes per module were determined based on the ranked kME order per Module. The grey module was disclosed from further analysis, since it contained arbitrary connected ncRNAs. Trajectories for each module were determined drawing the k-Module Eigengene vector for each timepoint and sample.

### NOCICEPTRA Tool and data sharing

To comply with FAIR and open access policies, we updated our recently published NOCICEPTRA Tool with the data obtained from this analysis. This tool will be made available open source at GitHub as an updated version of NOCIEPTRA, which also includes the necessary requirements for self-deployment using Python Streamlit. In addition, the tool will be made available online at https://nociceptra.streamlit.app and as source code at Github (muiphysiologie/NOCICEPTRA2.0). Furthermore, small ncRNA data and long RNA data obtained from hDRGs will be available dbGap (Accession number: phs001158). The small RNA data for iDN analysis is available from GEO Datasets (Accession ID: GSE161530). In addition, all analysis Python and R-scripts as well as data pipelines will be made available in a separate GitHub repository mouzkolit/Nociceptra2_Analysis (Zeidler *et al*. 2021).

## Results

### Time dependency of small RNAs throughout human iPSC-derived sensory neuron differentiation

In this study, we report on trajectories and hub ncRNAs to provide the first ncRNA atlas throughout the development of human iPSC-derived nociceptors (iDNs). Since our recent analysis (Zeidler *et al*. 2021) of mRNA and miRNA interactions excluded other relevant ncRNA species, we now extended the analysis towards tRNA, snoRNA, rRNA, lncRNA, piRNA and lincRNA expression patterns during iDN development from three different iPSC clones. To retrieve counts of different ncRNA species, trimmed small RNA sequencing reads were counted and aligned using the recently published eXCerpt pipeline against the human genome (hg38), miRBase.org, gtRNAdb and piRNAdb database in the eXCerpt pipeline’s default order. General raw count frequency analysis across samples revealed that the majority of ncRNA reads were annotated as miRNAs, followed by tRNAs, miscRNAs, snoRNAs and further mapped to other annotations such as intron retentions and protein-coding genes (**Supplementary Figure S2 A**, (Chan & Lowe 2009, 2016; Consortium *et al*. 2020; Piuco & Galante 2021)). In addition, distributional frequencies per biotype are also mostly conserved across samples and clones within a timepoint (**Supplementary Figure S2 A**). Aggregation on timepoint revealed that miRNAs exhibited low distributional changes in the frequency of counts throughout differentiation, where only day 26 showed a ∼10% reduction in the relative number of counts compared to all other timepoints (**Figure 1A**).

**Figure 1:**
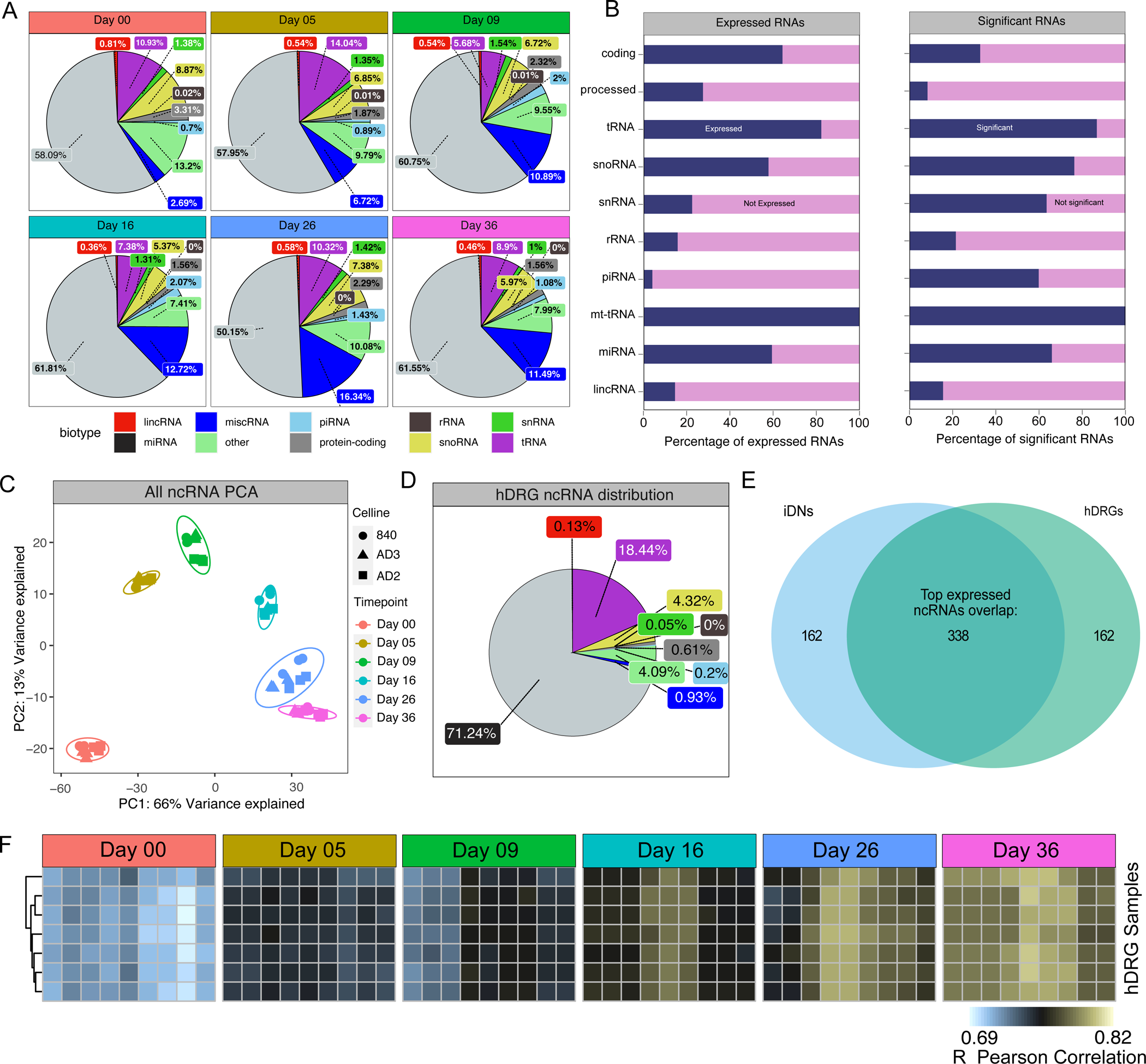
Basic exploratory analysis of ncRNA reads aligned using the exCerpt pipeline. **(B)** Timepoint and ncRNA biotype aggregated raw count frequencies were calculated using the average of each biotype per Timepoint. **(B)** Stacked-bar chart showing percentage of expressed vs unexpressed genes (below 5 counts, left panel), and the percentage of significant genes from the subset of expressed genes as indicated by the DESeq2 analysis (right panel). **(C)** Principal Component Analysis of variance stabilized counts were used to categorize samples into cell line and timepoint specific shapes and colors. (D) Distributional analysis of count frequencies aggregated based on ncRNA biotype for hDRG. (E) Set intersection analysis of the top 500 ncRNAs expressed in late-stage nociceptor (Day26-36) and hDRG. (F) Correlation analysis using Pearson correlation of variance stabilized counts between hDRG samples and iDN at the different timepoints revealed strong time -dependent approximation between hDRG and iDNs ncRNA signatures.

In contrast, tRNAs and snoRNAs exhibited pronounced timepoint dependent frequencies, and tRNAs accounted for 10.93% and 14.04% of raw counts in the iPSC-stage and early differentiation phases. Lower frequencies were identified at D09 (5.68%) and D16 (7.38%), and a small increase in tRNA count frequency was observed in more mature sensory neurons at D26 and D36 (**Figure 1A**). While snoRNAs showed a general decline in frequency throughout sensory neuron development, with 8.68% of counts annotated at D0 and only 5.97% of counts annotated at D36, miscRNA (miscellaneous RNA) showed anti-correlated patterns with low frequencies in the iPSC-stage (2.69%) and high number of counts at D26 (16.34%) and D36 (11.49%, **Figure 1A**). The majority of annotated tRNAs, rRNAs, snoRNAs, mt-tRNAs and miRNAs maintained at least low expression levels throughout development (**Figure 1B**). Differential gene expression analysis using the model ∼clone*timepoint (reduced model: ∼clone) revealed that the majority of expressed ncRNAs were differentially expressed (8000 aligned ncRNAs; 3000 with a base mean above 20) throughout development at any of the timepoints (**Figure 1B, right panel**). To evaluate the most significantly changed ncRNAs, differentially expressed ncRNAs were ranked based on the negative log10 adjusted p-value. In pluripotent iPSCs (D00), the lincRNAs CTD-244A18.1 and LINC00678, the tRNA-Thr-AGt-3 and SNORD90 were enriched. The piRNAs hsa_piR_018165 and hsa_piR_016659 were highly enriched at D16 (**Supplementary Figure S1 A**). Interestingly several snoRNA also exhibited abundant expression at D16 such as SNORA80B and SNORA73B with currently unknown functions for neuron development (**Supplementary Figure 1A**). As most regulated ncRNAs, tRNA-Glu-CTC-1 (upregulated at D26-D36), tRNA-Gly-GCC-1 (upregulated at D5) and hsa_piR_016658 (upregulated at D16) were highly abundant and significantly expressed throughout development (**supplementary Figure S1 B**).

To further evaluate the general time dependency of ncRNAs throughout development, principal component analysis (PCA) revealed a high degree of variance (PC1, 66% variance ratio) explained by the different time-points, which implicates pronounced ncRNA changes throughout the development of iDNs (**Figure 1C**). In contrast, much smaller clonal variances were detected, indicating that developmental changes were much more drastic than clonal differences (**Figure 1C**). This supports the idea that conserved biological ncRNA expression signatures may serve specific developmental stages during neuron differentiation and maturation.

In addition, hDRG samples from seven organ donors were analyzed, and distributional analysis revealed inter-sample variance of ncRNA biotypes, where miRNAs (71.24%) and tRNAs (18.44%) were the predominant expressed biotypes, followed by snoRNAs (4.32%) and snRNAs (4.09%) (**Figure 1D**). However, the pronounced distributional variance between the samples might be correlated to donor metadata information, but currently the sample-size is too low to infer confounding donor covariates (**supplementary Figure S2 B**) We used correlation analysis of normalized, variance stabilized counts to infer how similar iDN samples were to hDRG samples at different timepoints. iDN ncRNAs signatures became increasingly correlated with hDRG samples throughout the differentiation process (Day 00 R = ∼0.69, Day36 R = ∼0.84) (**Figure 1F**). Although inter-sample variance of hDRG samples was relatively high, correlation analysis with iDN samples revealed conserved similarities scores across all hDRG samples indicating similar intra-sample ncRNA expression signatures between hDRG samples (**supplementary Figure S2 C-D)**. hDRG and late-stage iDNs shared several top expressed miRNAs such as the let-7 family which were highest expressed in hDRG, and miR-182/183, which was the top expressed miRNAs in iDNs (**Supplementary Figure S3 A-B**). Trajectories of the let-7 family showed a strong upregulation towards more mature iDNs (> Day26, **supplementary Figure S2 F**) supporting a critical role of the let-7 family in maturation. miR-21, miR-100, miR-30d and miR-30a were consistently found amongst the list of top-20 expressed miRNAs in hDRG and all developmental iDN stages (**supplementary Figure S3 A-B**). ncRNA signatures differences were likely explained by the fact that our hDRG samples not only contained sensory neuron but also other cell-types such as Schwann cells, satellite cells and immune cells whose ncRNA profiles differ considerably from neuronal ones.

### ncRNA modules with preserved trajectories

To classify ncRNAs into highly interconnect modules representing co-expressed units, we filtered ncRNAs (baseMean > 20 as a metric of abundance and to remove spurious correlations) and used weighted gene correlation network analysis (WGCNA). This revealed 14 modules of highly correlated ncRNAs, and allowed us to identify modules with conserved expression trajectories found in all three iPSC clones such as the black, yellow, green and blue module. Conversely, we also detected modules associated with expression changes of ncRNAs found only in one of the iPSC-clones indicating clonal effects (e.g. magenta and red module) (**Figure 2 A-B**). Module trajectories of correlated ncRNAs were obtained using eigengene vector calculations (WGCNA R package) and associated with all stages of iDN development as defined previously (Zeidler *et al*. 2021), **Figure 2A**). Especially, the black and yellow modules were associated with the iPSC- and early differentiation stages and hub ncRNAs were classified as tRNAs such as tRNA-Thr-AGT-3 and tRNA-Val-TAC-4 (both black module), tRNA-MET-CAT-chr6 and tRNA-Lys-TTT-7 (both yellow module), as well as snoRNAs such as SNORD88B (yellow module), and lincRNAs such as RP3-467D16.3 (black module) and MIR17HG (yellow module). In contrast, hub snoRNAs, such as SNORD90, SNOR24, SNORD116 and SNORD32A, associated with the green and blue modules, were highly up-regulated during later stages (D16-D36). Of these, SNORD116 variants are known as important regulators of neuronal differentiation (**Figure 2B**, (Pace *et al*. 2020)). Interestingly, the majority of ncRNAs associated with the brown module were annotated as Y RNAs (Fragments). These Y RNAs such as Y_RNA.439 and Y_RNA.700 were highly enriched from D05-D16 implicating putative functions in neural crest cells and progenitor cell development into a sensory neuron fate (**Figure 2B**).

**Figure 2:**
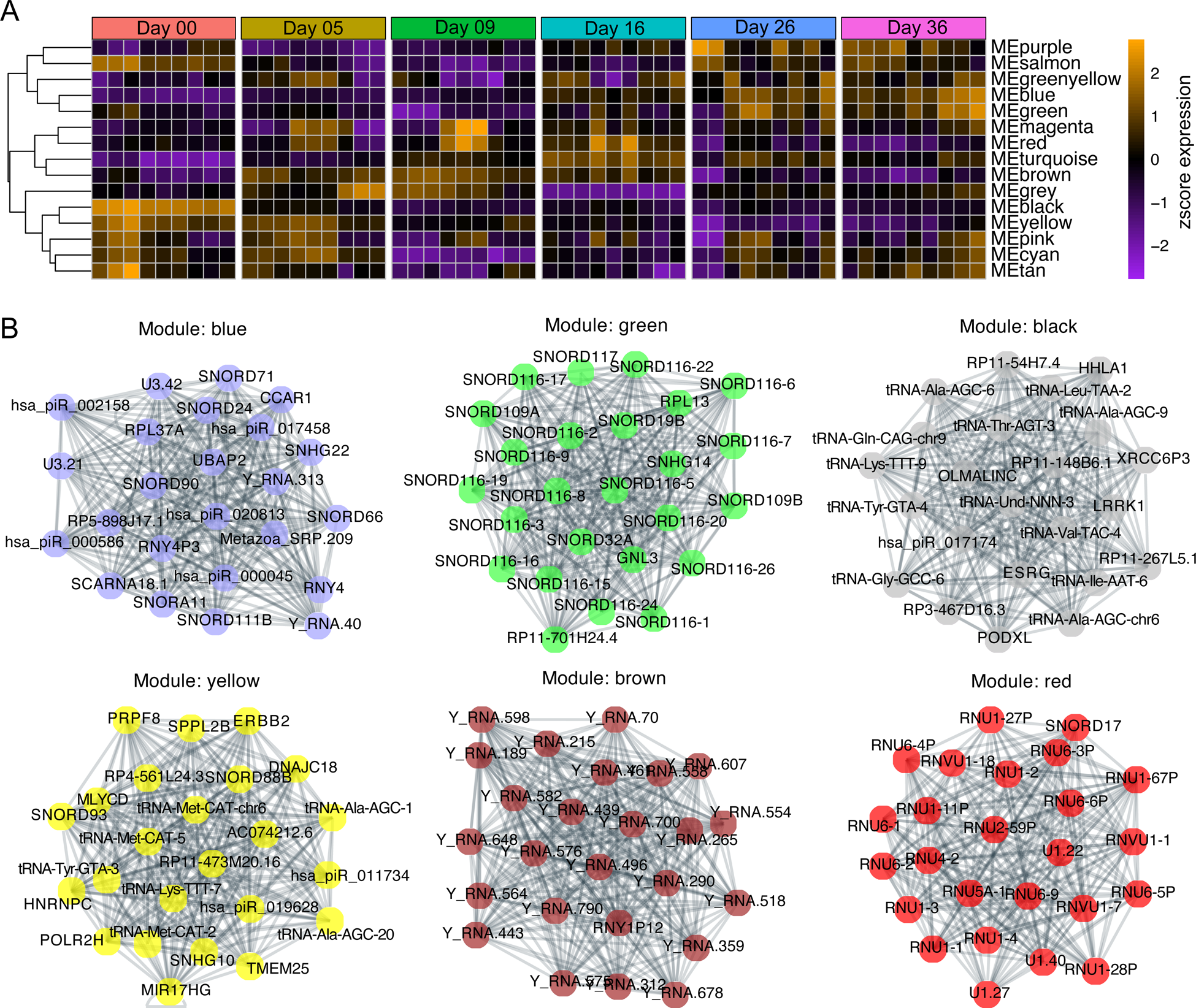
WGCNA analysis of ncRNA variance stabilized counts excluding miRNAs and only using ncRNAs with a base mean count above 20. **(A)** Eigengene vector trajectories of WGCNA derived modules were clustered using soft-threshold correlation into classes exhibiting the same temporal signatures. **(B)** Top 20 ncRNA hub gene network determined using determining the Pearson correlation coefficient between all module members using variance stabilized counts. Networks were drawn using the NetworkX python module using the correlation between ncRNA nodes as weight and the Netgraph module with the spring layout and the correlation score as edges weight.

### Differential temporal distribution of tRNA fragments during sensory neuron development

Transfer RNAs (tRNAs) represent a well-known ncRNA species serving RNA translation into protein, however, novel regulatory functions for tRNA fragments (tRFs) have recently been reported (Anderson & Ivanov 2014; Magee & Rigoutsos 2020). tRFs are short, 16-35 nucleotides in length, and play vital roles in neuron survival, necrosis, and neurodegenerative disorders such as Parkinson and Alzheimer’s disease, but have not been explored in neuronal development (Cao *et al*. 2021; Inoue *et al*. 2020; Mathew *et al*. 2022; Schaffer *et al*. 2019; Wu *et al*. 2021). tRNAs and tRFs exhibited pronounced time dependent expression patterns throughout iDN differentiation supporting the idea that tRNA regulatory pathways contribute to neurodevelopmental processes (**Figure 1A**). A more in-depth analysis of count frequency for amino acid specific tRNAs revealed that tRNA^Gly^ and tRNA^Glu^ were the most frequent tRNA fragments throughout differentiation with inversely correlated frequency patterns: while tRNA^Gly^ was more abundant early in development, tRNA^Glu^ predominated in transition stages such as D09 and was highly abundant in late iDN stages (**Figure 3A**). Most counts (average read length ∼25 nucleotides long) emerging from the tRAX pipeline represented tRFs rather than full length tRNAs, which may partially result from potential limitations of the underlying sequencing technology (**supplementary Figure 4A**) (Holmes *et al*. 2022). However, the percentages of tRNA^Lys^ (7.94%), tRNA^Asp^ (6.94%) and tRNA^His^ fragment counts were strongly increased at D09 in comparison to all other days, indicating a putative shift not only in individual tRNA species but also tRNA fragment types and this finding supports specific regulatory roles of tRFs to control gene sets required for pluripotency vs. neuron differentiation and maturation.

**Figure 3:**
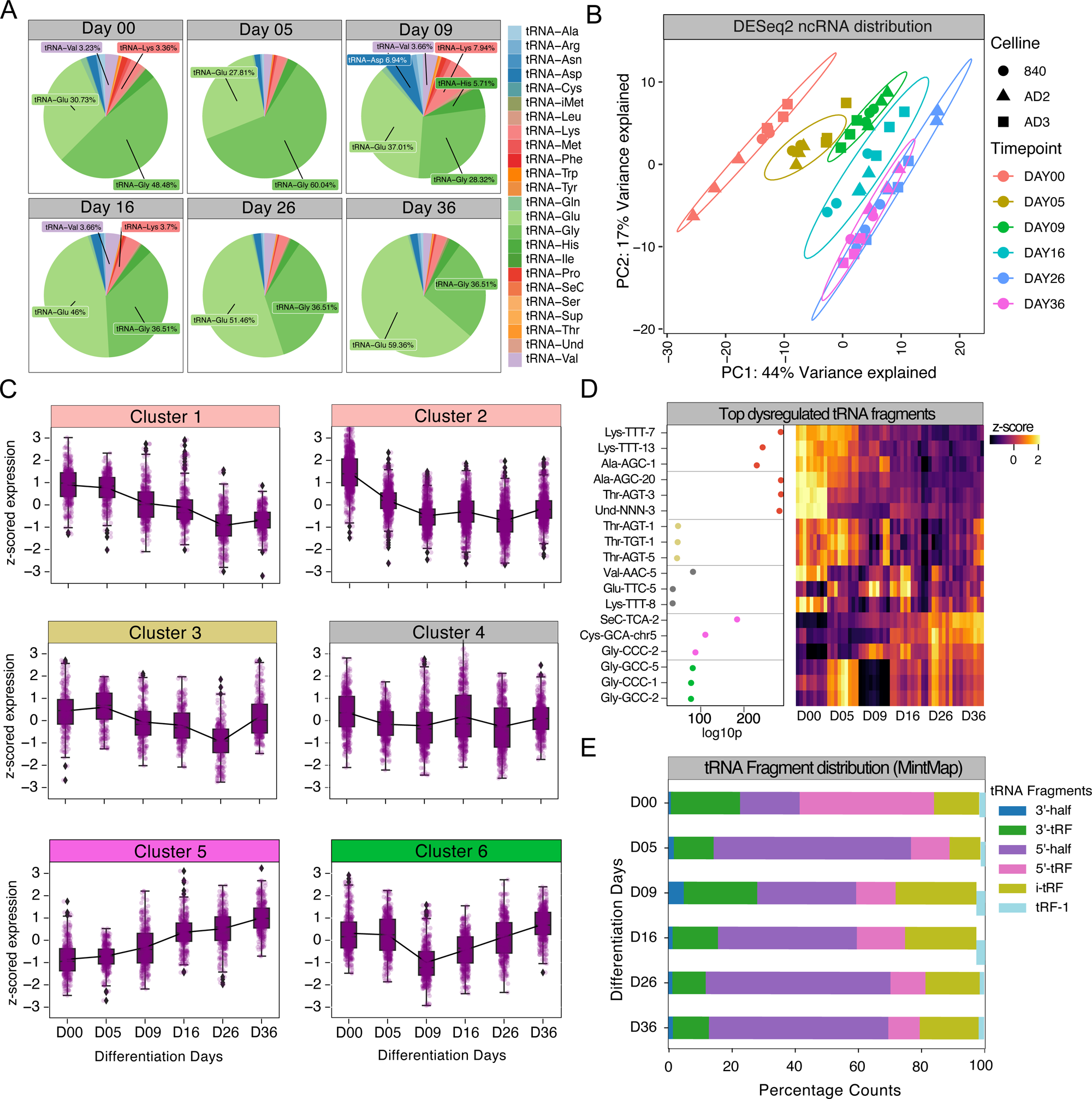
tRNA specific analysis of developmental trajectories. **(A)** tRNA specific raw counts were aggregated on timepoint and tRNA amino acids and count distribution was depicted as pie chart. **(B)** PCA of tRNA variance stabilized counts was clustered based on cell line and timepoints. **(C)** Hierarchical cluster analysis using the distance metric (1-correlation) as distance and the ward algorithm to determine clusters was used, which revealed 6 clusters of unique tRNA expression trajectories. Trajectories are represented as z-scored scaled variance stabilized counts and both aggregated counts (boxplots) and unique counts are shown. **(D)** Top regulated tRNAs per cluster are depicted by means of the negative log10 p-adjusted value as well as the expression patterns. **(E)** Sequencing reads were further aligned using MintMap2, a specific pipeline to count tRNA reads and determine location of tRNA fragments, while also revealing specific sequence of ncRNA contigs as well as the specific fragment type. Fragment types per timepoint were aggregated using the sum function within samples and the average function across samples from the same timepoint.

To explore the dependencies between neuronal development and tRNA expression signatures two approaches were used: first, only tRFs determined by eXcerpt were used for analysis (**Figure 3B-D**); second, tRFs were realigned using tRNA specific tools such as MintMap2 to reclassify tRNA fragments based on mapping occurrence (**Figure 3E**). eXcerpt normalized counts of annotated tRNAs followed by PCA revealed pronounced time dependency for tRNA fragments (**Figure 3B**). To retrieve information about co-expressed tRFs, all expressed tRNA sequences were subject to hierarchical clustering which revealed six stages characterized by tRF expression during sensory neuron differentiation (**Figure 3C**). By ranking of each cluster based on the FDR-corrected p-value and plotting of trajectories we found top-regulated tRFs for each cluster such as tRNA-Lys-TTT-9-1 (iPSC stage, early differentiation, cluster 1), tRNA-SeC-TCA-2 (nociceptor stage, cluster 4), tRNA-Gly-GCC-2 (nociceptor, cluster 5) and tRNA-Gly-CCC-5-1 (early differentiation phase/nociceptor, cluster 6, **Figure 3D**). In addition, mintMap2 was used to classify the exact fragment type of the exclusive tRF spaces, and distributional analysis throughout iDN differentiation was performed (Rozowsky *et al*. 2019). Most reads classified as 5’ tRFs (5’ halves, 5’tRF, as supported by the tRAX analysis) followed by 3’ tRFs (3’ halves, 3’ tRFs) throughout differentiation, but also showed proportional changes between 5’halves and 5’tRFs. While 5’tRFs were enriched in iPSCs, 5’ halves became more abundant throughout differentiation, which might indicate differential cleavage of tRNA fragments throughout development. In addition, i-tRFs (internal tRNA fragments) showed the highest compositional fraction at D09 (**Figure 3E**). To further evaluated putative roles of the most significantly tRNA fragments per cluster, we used the DIANA MR-microT software and queried the aligned sequences derived from the MintMap2 alignment for putative target mRNA spaces. The tRNA GlyCCC was likely targeting genes implicated in Axon Guidance (KEG:04360) and GluCTC appeared to regulate cell projections and synapse related pathways (**Supplementary Table S5**). Interestingly, both tRNA-GluCTC as well as tRNA-GlyCCC significantly decreased at Day 09, which is considered the starting point of extensive axon growth. This might support the idea that these two tRNA fragments may play a significant role in axonogenesis/synaptogenesis (**Supplementary Figure 5A**). In contrast, ThrAGT and ThrTGT were likely involved in RNA transcription and metabolic process regulation (**Supplementary Table S5**).

We also evaluated tRNAs likely mimicking miRNAs and queried tRNA targeted genes for each individual tRNA fragment to retrieve miRNA interactions using microTCDS v.5. GlyCTC and GlyCCC mimicked the functionality of the miR-548 family and CycACA and AlaAGC shared target spaces with miR-3163, miR-4282 and miR-4536-5p (**Supplementary Figure S5 B**). In addition, overlap analysis of hDRG and iDNs revealed that a significant portion of the top-20 expressed tRNA fragments were found both in hDRG and iDNs such as member of the tRNA-GlyGCC, tRNA-GluCTC, tRNA-AspGTC and tRNA-LysCTT family (Supplementary Figure S3 A-B).

Finally, we evaluated expression changes of mitochondrial (mt-) tRNAs throughout differentiation and found that all mt-tRNAs were differentially expressed with increasing expression throughout sensory neuron development, which putatively correlated with an increased expression and translation of mitochondrial proteins to secure the increasing energy requirements of maturating neurons (**supplementary Figure S4** (Cao *et al*. 2020; Suzuki *et al*. 2011)).

### Differential temporal distribution of snoRNAs in sensory neuron development

snoRNAs are well accepted for their role in ribosomal protein regulation but expression signatures and regulatory roles in human neuronal development are mostly unknown (Huang *et al*. 2022; Kufel & Grzechnik 2018). PCA on all expressed snoRNAs revealed high temporal dependencies of snoRNAs indicating a strong regulatory effect of snoRNA expression throughout differentiation (**Figure 4A**). Six clusters emerged with differential temporal snoRNA expression signatures associated with different developmental stages, with cluster 1 and 2 representing the late stages of development, stage 3 the progenitor stage/immature neuron stage, clusters 4 and 5 the early developmental stages and cluster 6 which represented snoRNAs downregulated in iPSCs (**Figure 4 B**). SNORD93 (baseMean: 1618 normalized counts), SNORA69.1 (baseMean: 37 normalized counts) and SNORD88B (baseMean: 74 normalized counts) were the most significant differentially expressed snoRNAs associated with the iPSC or early development stages while SNORA80B (base-mean: 25 normalized counts), SNORA73B (base-mean: 731 normalized counts) and SNORD94 (base-mean: 281 normalized counts) were significantly associated with D16. Their roles and the functional consequences of this regulation are currently unknown. SNORD71 (baseMean: 8436), SNORD115 (baseMean: 40 normalized counts) and SNORD116 (baseMean:1629 normalized counts) were up-regulated in more mature neuron-like cells (**Figure 4B-C**). Of these, SNORD115 promotes serotonin receptor (HTR2C) activity and for SNORD116 roles in neuron development, differentiation and survival are already known (Bratkovič *et al*. 2018; Burnett *et al*. 2017; Pace *et al*. 2020; Polex-Wolf *et al*. 2018). The precise function and regulated pathways are currently unknown for the majority of snoRNAs such as for abundantly expressed SNORD71, SNORD89 and SNORD90, which both exhibited a gradual increase in expression throughout differentiation (**Figure 4 C**) indicating that specific regulatory roles may apply to other snoRNAs as well. This is further supported by our finding that the majority of the top 20 expressed snoRNAs were conserved between hDRG and iDNs such as SNORD30, SNORD104, SNORD48 and SNORD69. Since snoRNAs modifying RNAs via methylation (C/D) (C/D Box snoRNAs) are distinct from snoRNAs which introduce pseudouridination (H/ACA) (H/ACA snoRNAs) we investigated the distribution of the two snoRNA classes throughout development (**Figure 4 D**). Significantly regulated C/D box snoRNAs (cluster 1, 2) were enriched throughout late differentiation (**Figure 4 D**), while expression H/ACA snoRNAs was higher in the early stages of iDN development (Cluster 5). Since functional consequences of snoRNAs are difficult to depict we focused further on the expression of host-genes (gene that is harboring the ncRNA) associated with snoRNAs and correlated host-gene with snoRNA expression. Significantly correlated and anti-correlated host-genes with R > abs[0.7] were subject to ontology enrichment analysis. This revealed detrimental changes in ribosomal regulation throughout sensory neuron development for both positively as well as negatively correlated snoRNA host-genes as indicated by the expression patterns of ribosomal host-genes (**supplementary Figure S4 B-C, supplementary Table S2/S3 positive-correlated/negative correlated**).

**Figure 4:**
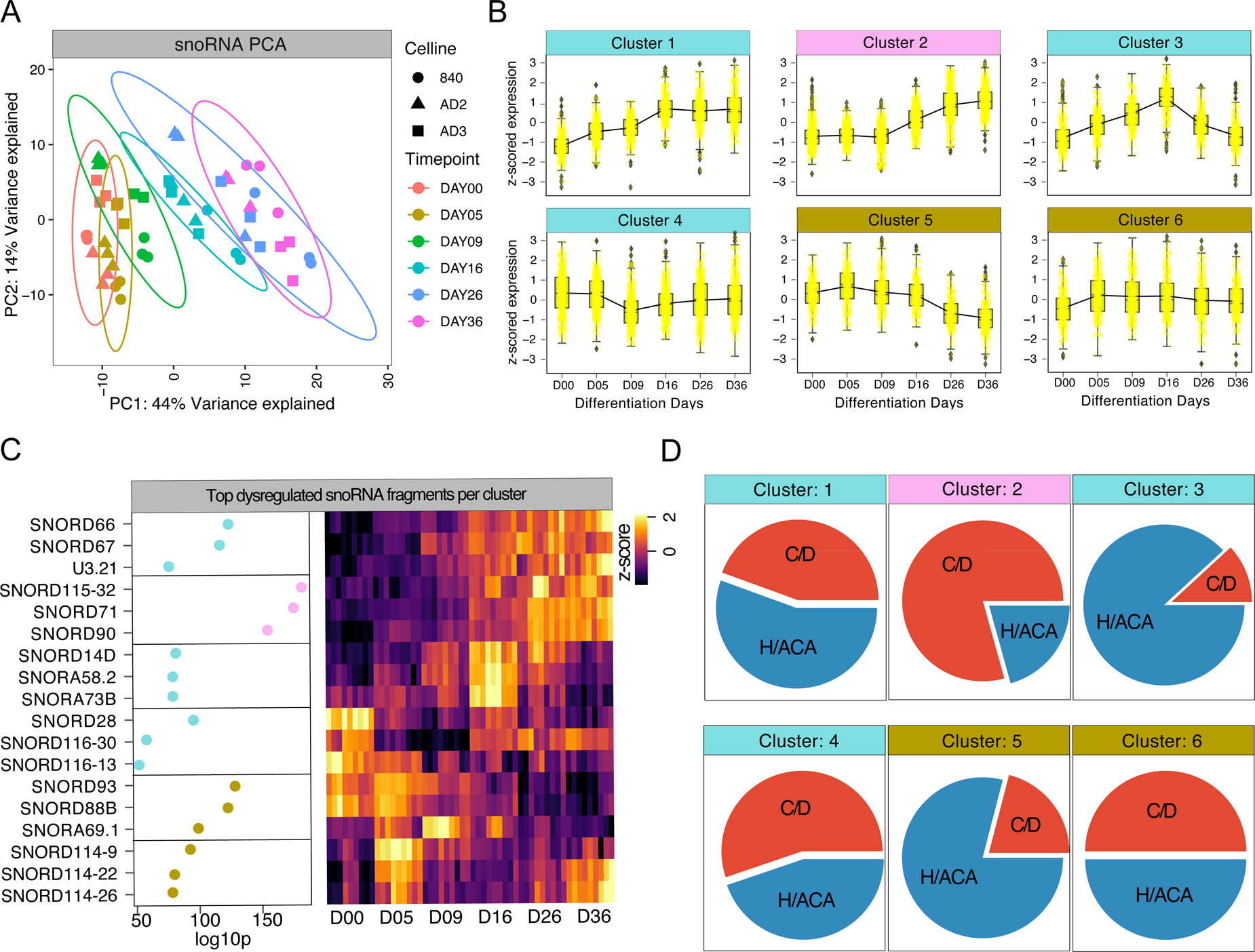
snoRNA specific analysis of development trajectories. **(A)** PCA of variance stabilized snoRNA counts revealed separation of timepoints. **(B)** Hierarchical clustering (using the distance 1-correlation) analysis grouped snoRNA into 6 cluster describing iDN development **(C)** Ranked top regulated members based on the p-adjusted FDR corrected p-value of snoRNAs per cluster and expression signatures were represented as z-score variance stabilized counts using a heatmap. **(D)** snoRNA class affiliation was determined using the snoDB database. Number of snoRNAs members per group (C/D, H/ACA) were determined per timepoint summing counts per each sample individually and average across the hierarchical cluster.

### Temporal regulation of rRNAs, snRNAs and piRNAs

We also evaluated the temporal trajectories of ribosomal RNAs (rRNA), small nuclear RNAs and piwi-interacting RNAs (piRNAs) using PCA. snRNAs and rRNAs showed only minor time-dependent changes throughout nociceptor development (**Figure 5 A-B**). Nonetheless, clustering analysis revealed five clusters associated with snRNA and three clusters associated with rRNA expression (**supplementary Figure S6**). Clonally conserved RNU5E-6p and RNU7-3P ncRNAs were highly expressed early in development (associated with cluster 1) and decreased with increasing differentiation and maturation indicative for a regulatory role in suppressing pluripotency genes. RNU2-3P, RNU5E-1 and RNU7-19P expression increased from Day 09 to Day 36 (cluster 1, 3, **supplementary Figure S4 B**). In contrast, piRNA fragments with a usual length of 21-24 nucleotides exhibited pronounced temporal dependency and clustering based on timepoint associated with sensory neuron development (**Figure 5C**). Small silencing piRNAs are implicated in the silencing process of transposable elements in germ-line cells thereby ensuring germline DNA integrity (Iwasaki *et al*. 2015) Although the cumulative number of piRNA reads was low, hierarchical clustering revealed five well distinguishable clusters of piRNA expression signatures (**Figure 1B, Figure 5D-E**). Cluster 1 and 2 piRNAs with the most significant members hsa_piR_011968, hsa_piR_019628 exhibited abundant expression in the iPSC stage and early during development. Cluster 3 piRNAs (hsa_piR_018165) were upregulated at D09-D16 and cluster 4-5 piRNAs (hsa_piR_00170, hsa_piR_002158) showed abundant expression at late stages (D26-D36, **Figure 5D-E**). Interestingly, the overall percentage of piRNA counts was considerably lower in hDRGs compared to iPSC (0.2% in hDRGs and ∼1% in iDNs). However, the highest expressed piRNAs showed a strong overlap between hDRG and iDNs, and the highest expressed piR-016658, piR-017033 and piR-018780 were conserved (**Supplementary Figure S3 A-B**). Based on the specific expression patterns and their trajectories, regulatory roles in neurodevelopment can be anticipated also for piRNAs.

**Figure 5:**
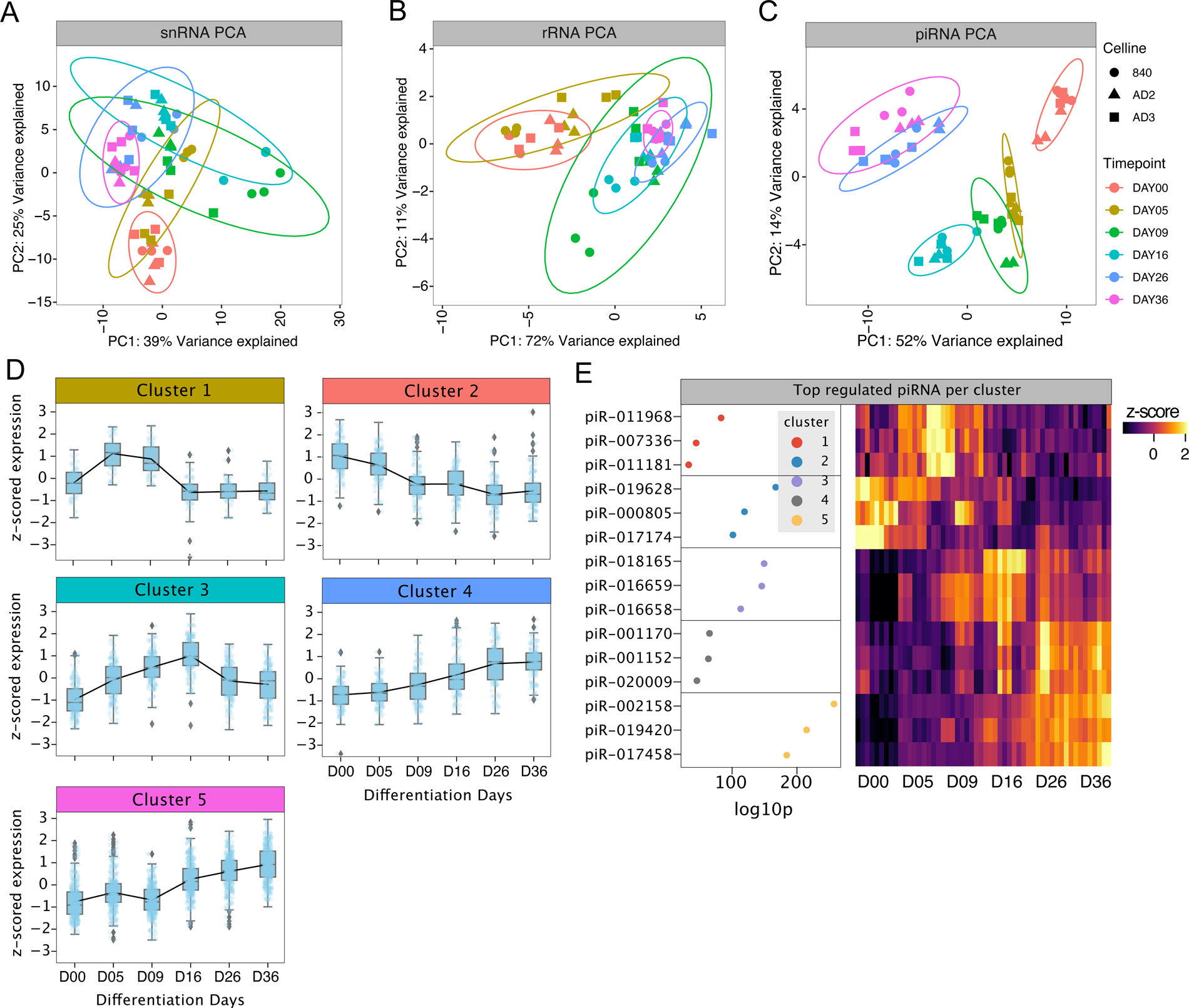
Analysis of snRNA, rRNA and piRNA trajectories derived from variance stabilized counts. **(A)** PCA of snRNA variance stabilized counts show low temporal dependency as depicted by the distribution of samples and the variance explained. **(B)** PCA of rRNA although exhibit low temporal dependencies based on the variance explained. **(C)** PCA of piRNA show high temporal dependency and timepoint dependent clustering. **(D)** Hierarchical clustering (distance: 1-correlation) analysis of z-scored variance stabilized piRNA counts revealed 5 distinguishable clusters. (E) Top ranked dysregulated piRNAs per cluster visualized by the negative log10 p-value and expression signatures are represented by the z-scored standardized variance stabilized counts.

### Long non-coding RNAs

We further investigated long non-coding RNAs with and without poly-A tails by integrating both the mRNA with the small ncRNA sequencing dataset. lincRNAs derived from small RNA sequencing and lncRNAs from long RNA sequencing exhibited pronounced temporal dependencies during iPSC differentiation into sensory neurons (**Figure 6A, B**). Clustering analysis revealed 5 lincRNA groups and 7 lncRNA groups with distinct temporal signatures (**Figure 6C, supplementary Figure 4 A**). lncRNAs B3GALT5-AS1 (ENsG00000184809) and LINC01356 (ENSG00000215866) were highly enriched in the iPSC-stage. In contrast, LINC00205 (ENSG00000223768), and FTX (ENSG00000230590) exhibited signatures of nociceptor stage-related gene expression. (Li *et al*. 2019b) supports an important role of FTX in the survival and possibly function of mature human iPSC-derived nociceptors. In addition, SOX1-OT (ENSG00000224243) and RENO1 (ENSG00000287431) were highly abundant in cluster 2 (∼Day05-Day09). Both lncRNAs are functionally interesting since SOX1-OT promotes SOX1-dependent neuronal differentiation in human embryonic stem cells (hESCs) and RENO1 regulates neuronal commitment in mouse embryonic stem cells (Ahmad *et al*. 2017; Hezroni *et al*. 2020; Li *et al*. 2021). Based on these reports and our current findings functional relevance is anticipated for SOX1-OT and RENO1 trajectories for iDN fate decision and differentiation. Since overall count numbers for lincRNAs were low, we merged lincRNA and lncRNA clusters for further downstream analysis. In general, lncRNAs/lincRNAs act as sponges for miRNAs or other RNA fragments in neurons (Yao & Yu 2019). Therefore, we specifically searched for enriched miRNAs that putatively targeted lncRNAs per cluster. lncRNAs associated with pluripotency were enriched for miRNAs of the let-7b family, which is increasingly expressed throughout development and essential for neural stem cell differentiation and cell fate commitment (Deng *et al*. 2020; Zeidler *et al*. 2021; Zhao *et al*. 2010). lncRNAs abundant at D00-D05 were enriched for hsa-miR-16-5p, hsa-miR-15a|b-5p and hsa-miR-434, while at the mature nociceptor stage (D36) enriched lncRNAs targeted hsa-miR-574-5p, hsa-miR-485-5p, hsa-miR-369-3p and hsa-miR-143-3p. Especially lncRNAs enriched at D00 and D05 showed highly anti-correlated trajectories with their enriched miRNAs, suggesting putative regulatory controls of these miRNAs in pluripotent cells by interactions with the enriched lncRNAs or vice versa (**Figure 6D**).

**Figure 6:**
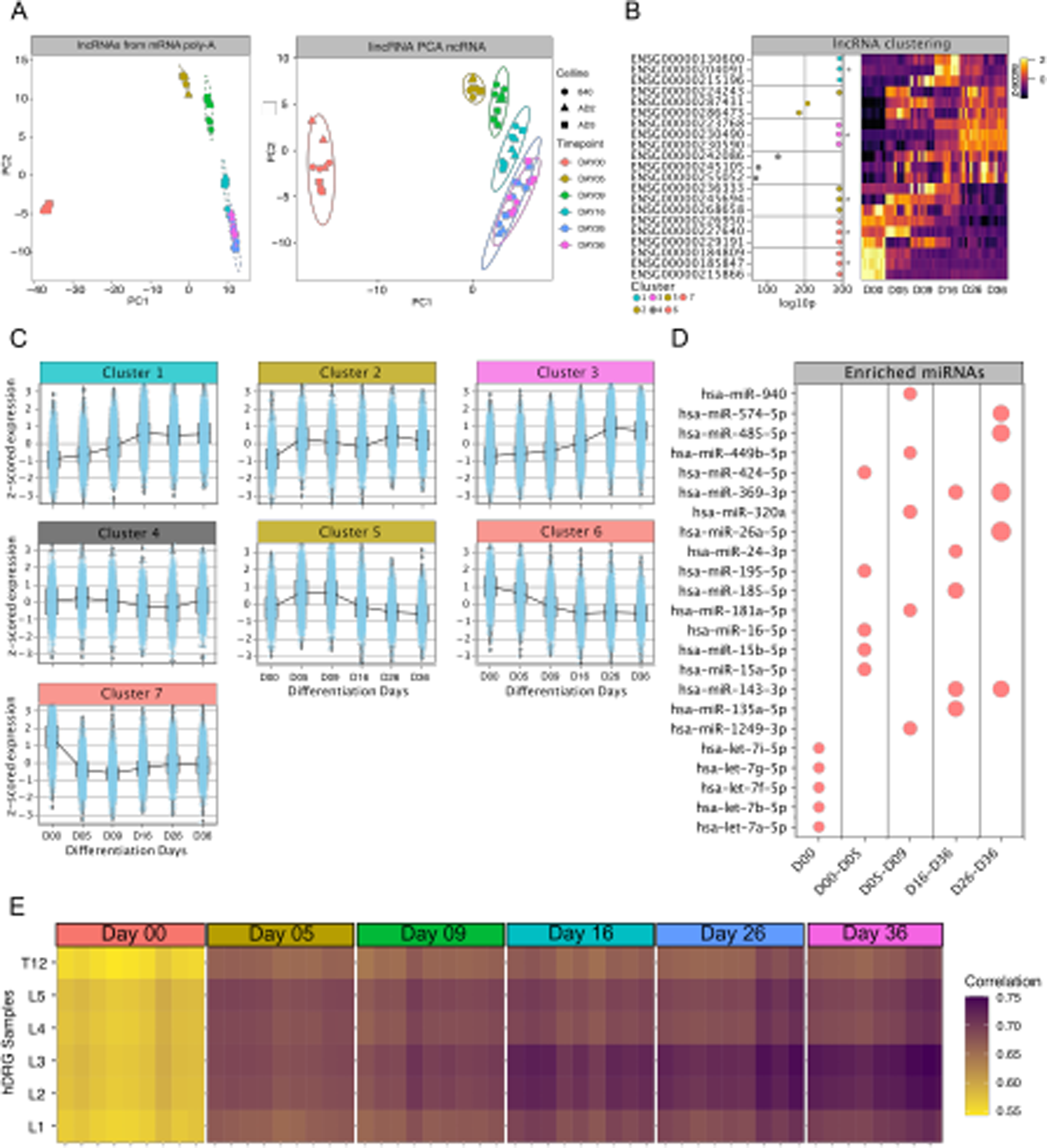
Analysis of lncRNAs and lincRNAs from long and small RNA sequencing reads. **(A)** PCA from lncRNA variance stabilized counts obtained from the mRNA sequencing. **(B)** PCA from lincRNAs obtained from small RNA sequencing without poly-A tail. **(C)** Clustering analysis of lncRNAs using hierarchical clustering revealed 7 cluster of lncRNA regulation throughout sensory neuron development. **(D)** lncRNA and lincRNAs clusters were merged into one cluster holding both groups, based on trajectories and assigned lincRNAs and lncRNAs were subjected to the lncSEA algorithm to determine potential enrichment of miRNAs per cluster. Enriched miRNAs per cluster were determined based on the number of lncRNAs interacting with the miRNA (dot-size). **(E)** Correlation analysis using variance stabilized counts of lncRNAs at all timepoints in iDN development and in hDRG samples revealed a significant increase of similarity in lncRNA expression throughout iDN development.

Finally, we compared iDN lncRNA expression signature to hDRG signature by means of a correlation analysis between all stages of iDN development and hDRG samples prepared from lumbar regions 1-5 and thoracal region 12. This revealed that maturating iDNs became more like hDRG neurons in terms of lncRNA expression signatures. Interestingly, iDNs were strongly correlated to neurons located in lumbar segment L2 and L3 hDRG whereas weaker correlation was identified for iPSCs (D0, **Figure 6E**).

## Discussion

While we previously determined miRNA trajectories and hub-miRNAs, this work extends the landscape of ncRNA regulation by revealing that ∼40-50% of the retrieved sequences are not classified as miRNAs but as other non-coding RNA species such as tRNA fragments, snoRNAs or miscRNAs in human iPSC-derived sensory neurons. Our distributional analysis of all currently known ncRNA species revealed conserved clonal temporal changes in count distribution at different timepoints throughout development for tRNAs, snoRNAs, piRNAs, lncRNAs, miscRNAs and lincRNAs. This putatively indicates that ncRNAs in general and not only miRNAs are functionally important and need to be considered as phenotypic cellular signatures of pluripotent cells and differentiating human nociceptors development in greater detail.

Several ncRNA species including miRNAs, snoRNAs and lncRNAs are emerging as important cell and tissue specific hub regulators of processes involving protein-coding genes and other RNA products (Guthrie & Patterson 1988; Iwasaki *et al*. 2015; Kumar *et al*. 2014; Winek *et al*. 2020). We carried out ncRNA transcriptomics from native human DRG, to provide insight in the distribution and expression signatures of different ncRNA biotypes and understand similarities and dissimilarities between native hDRG and iDNs. The combined analytical approach of iDN and hDRG, putatively helps to identify ncRNAs that are relevant for human sensory neuron differentiation and maturation. This is highly relevant, since in-depth analysis of ncRNAs is not possible through scRNA sequencing technologies yet. We developed an atlas for ncRNA expression of nociceptors reprogrammed from human iPSCs and revealed that additional ncRNA biotypes next to miRNAs contribute significantly to the developmental process in human stem cell derived neurons (Skreka *et al*. 2012; Stefani & Slack 2008). To enable researchers to re-use our data and investigate ncRNAs in depth, our significantly extended NOCICEPTRA2.0 tool will be provided locally at Github (https://github.com/MUIphysiologie/NOCICEPTRA2.0) or online via streamlit-cloud (https://nociceptra.streamlit.app) as an open-source web framework connected to the snowflake cloud, to enable easy accessibility to analyzed data. NOCICEPTRA2.0 is the first comprehensive ncRNA atlas to explore all currently known RNA species and their trajectories in detail, together with WGCNA modules for mRNA/miRNA and ncRNAs as well as indications of ncRNAs differentially expressed throughout development.

Five to seven developmental mRNA::miRNA clusters characterize specific stages of human nociceptor development (Zeidler *et al*. 2021), and these were also conserved for the majority of ncRNA biotypes. The majority of ncRNAs showed time-dependent expression trajectories throughout development, suggesting that specific ncRNAnome signatures are necessary to drive human iPSC-derived nociceptor development. Interestingly, few studies report detrimental changes in snoRNAs and tRNAs in differentiating mouse embryonic stem cells (Krishna *et al*. 2019; McCann *et al*. 2020). Our data confirmed increasing expression following differentiation initiation until D16 for Snora53, Snora54 and highly expressed tRNA-Gly-GCC early in development supporting important roles for Snora53/54 throughout the entire development process for neural progenitor cells whereas tRNA-Gly-GCC appeared to be involved in the general initiation process of differentiating neurons.

The majority of significantly regulated ncRNAs of different biotypes have not been reported or functionally associated with neuron development. This includes tRNA fragments such as tRNA-Thr-AGT-3 and tRNA-Ala-AGC-20, the snoRNAs SNORD71 and SNORD90, the piRNAs hsa_piR_019420 and hsa_piR_018165 and a whole group of miscRNA as most significantly regulated ncRNAs. tRNAs most likely represented tRNA fragments (cleaved tRNAs) while complete tRNA transcripts were rare probably due to the sequencing approach (Motorin *et al*. 2007). While the majority of tRNAs were annotated as 5’ and 3’ tRFs, differential cleavage throughout the development may account for shifting from tRFs to longer tRNA halves, indicating alterations in tRNA silencing and target spaces or a compensatory response to stress throughout development as tiRNA (tRNA halves) are upregulated in response to stress (Shen *et al*. 2018; Winek *et al*. 2020). In addition to the differential composition of fragment types throughout differentiation, we also showed that tRNA-isodecoder distribution shifted from glycine to glutamine throughout differentiation, which might indicate an increased usage of glutamine for translational processes in later stage iDNs.

Since it is well described that interactions of ncRNA fragments with mRNAs as well as target spaces are many-to-many relationships, we clustered all ncRNA biotypes into transcriptional units exhibiting the same expression signatures (Chipman & Pasquinelli 2019). Modules highly correlated with late-stage, more mature nociceptors consisting of multiple ncRNA biotypes such as SNORD90, which is enriched in brain tissues, orchestrating the development into iDNs (Isakova *et al*. 2020). In future it would be interesting to integrate these modules with temporal ATAC sequencing data to explore putative transcription factor binding sites correlated with the expression trajectories of ncRNAs (Li *et al*. 2018, 2019c).

In addition to the short ncRNAs, lncRNAs play important roles in neuron development by regulating cell fate determination, progenitor pool maintenance and proliferation, as well as neuronal survival (Li *et al*. 2019a, 2021). Non-surprisingly, lncRNAs exhibited the highest temporal to clonal variance ratio throughout iDN development, indicative of conserved roles for lncRNAs not only in neurodevelopmental processes of the brain (Li *et al*. 2019a; Ng *et al*. 2013; Rani *et al*. 2016; Sarangdhar *et al*. 2018) but also in human iDN and nociceptor development. In addition, lncRNA expression signatures of iDNs became more similar to hDRG lncRNA signatures throughout the development of iDNs suggesting that they predominantly originated from sensory neurons. This bolsters the suitability of the iDN model for in-vitro human sensory neuron research. Moreover, overexpression of those ncRNA that are highly expressed in hDRGs but lowly in earlier stages of differentiation of iDNs such as the let-7 family holds great potential as promising strategy to further enhance maturity and similarity of human iDNs and native nociceptors.

Evaluating the roles and contributions of specific ncRNA biotypes throughout iDN development revealed differential importance of ncRNA biotypes for the developmental process. While tRNAs, snoRNAs, piRNAs and lncRNAs exhibited pronounced temporal development specific dependencies, which putatively indicated critical contributions to development processes, snRNAs and rRNAs were only to a minor extent associated with the differentiation process in our analysis. Although this might be confounded by the overall shallow depth of the ncRNA sequencing approach, which only allowed to investigate more abundantly expressed ncRNAs. it could also support the idea towards a more general role of snRNA and rRNA regulation signatures conserved throughout all developmental stages. Changes in lncRNA, tRF and miRNA signatures with increasing maturation may also reflect increased electrical activity of more mature neuron and immediate changes in expression after activity, during which fluctuations in mRNA abundance subject to ncRNA regulation are observed (Barry *et al*. 2017).

In summary, developmental ncRNA expression signatures of iDNs were approximating signatures of native hDRG throughout development, and top expressed and regulated ncRNAs were surprisingly similar as indicated by our analysis. This strengthens the usage of iDNs for sensory neuron research, ad provides further opportunities to improve the maturity of iDNs. Overall, NOCICEPTRA2.0 provides the first comprehensive and online available, searchable atlas of currently known RNA species in developing and maturating iPSC derived human nociceptors which become increasingly important as human model systems for preclinical pain research.

## Funding

This study was funded by the Austrian Science Fund FWF (DK-SPIN, B16-06 to MK) and the European Commission (ncRNAPain, GA 602133 to MK).

## Acknowledgement

The authors thank C. Wild and M. Tschugg of the IT Services, Medical University of Innsbruck, for providing general support regarding the computing environment.

## Author contributions

MZ, TF and JB performed the experiments; MZ developed the NOCICEPTRA2.0 tool; MZ and MK conceptualized the study; MZ, TP and MK wrote the manuscript.

## Competing Interest

The authors declare no competing interests.

---

**Supplementary Figure S1:**
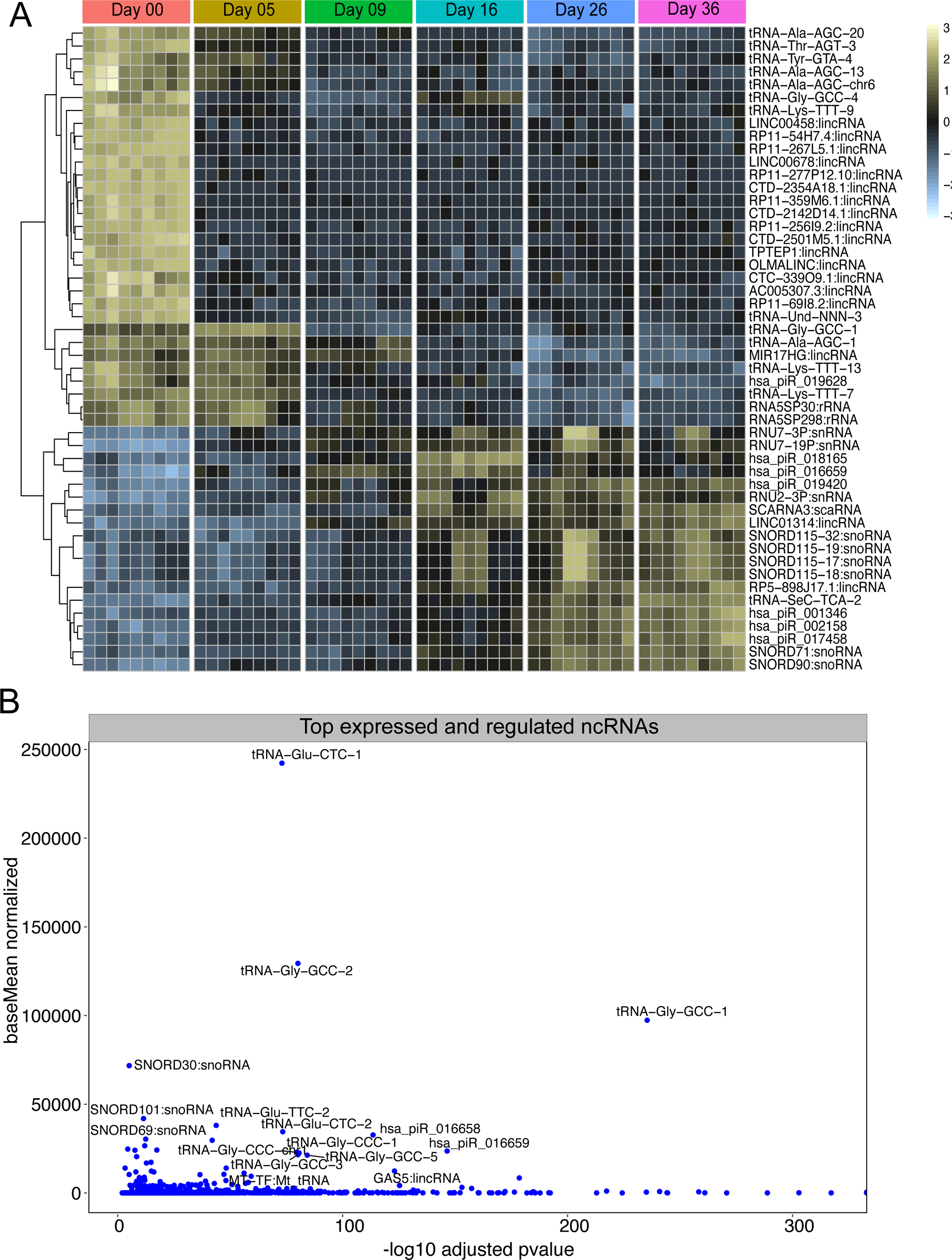
Differential gene expression analysis of all ncRNAs. **(A)** Top 50 regulated ncRNAs including snoRNAs, lincRNAs, tRNAs, piRNAs, snRNAs, scaRNAs clustered rows into same trajectories and split by differentiation timepoints (columns). **(B)** Highly expressed ncRNAs were depicted as a function of the negative log10 p-adjusted value to determined highly significantly expressed ncRNAs

**Supplementary Figure S2.**
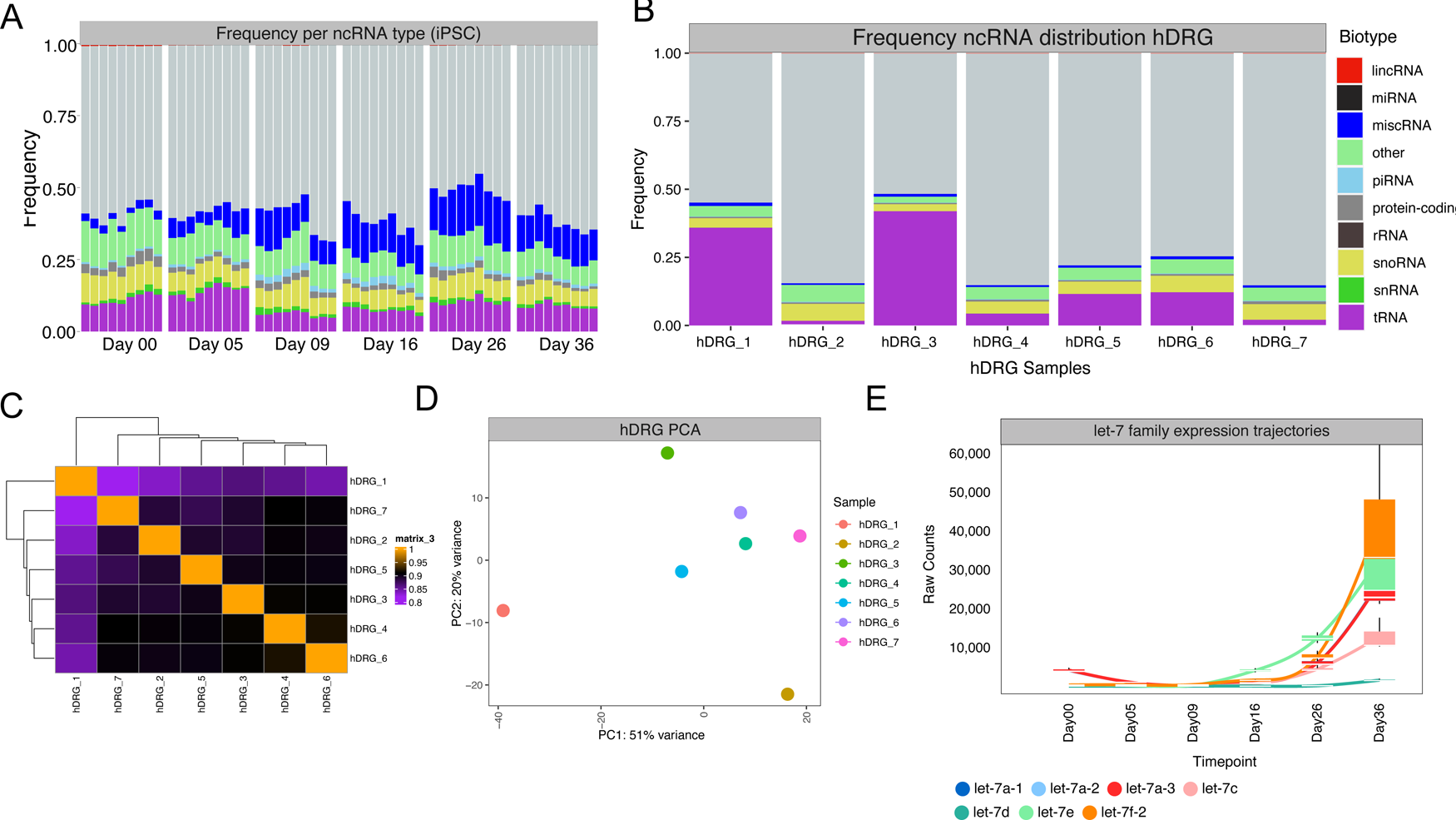
(A) iDN sample-wise distributional analysis of count frequencies aggregated on the biotype per sample (B) hDRG sample-wise count frequence aggregated per biotype/sample. (C) Hierachical clustering of the correlation dissimilarity of all hDRG samples (D) PCA of hDRG samples from variance stabilized counts. correlation between hDRG samples. Aggregated distributional analysis of ncRNAs per biotype using the sum aggregate function per sample and calculating the mean frequency per biotype across samples. (F) Expression trajectories of the hsa-let-7 family throughout development depicted using raw counts.

**Supplementary Figure S3.**
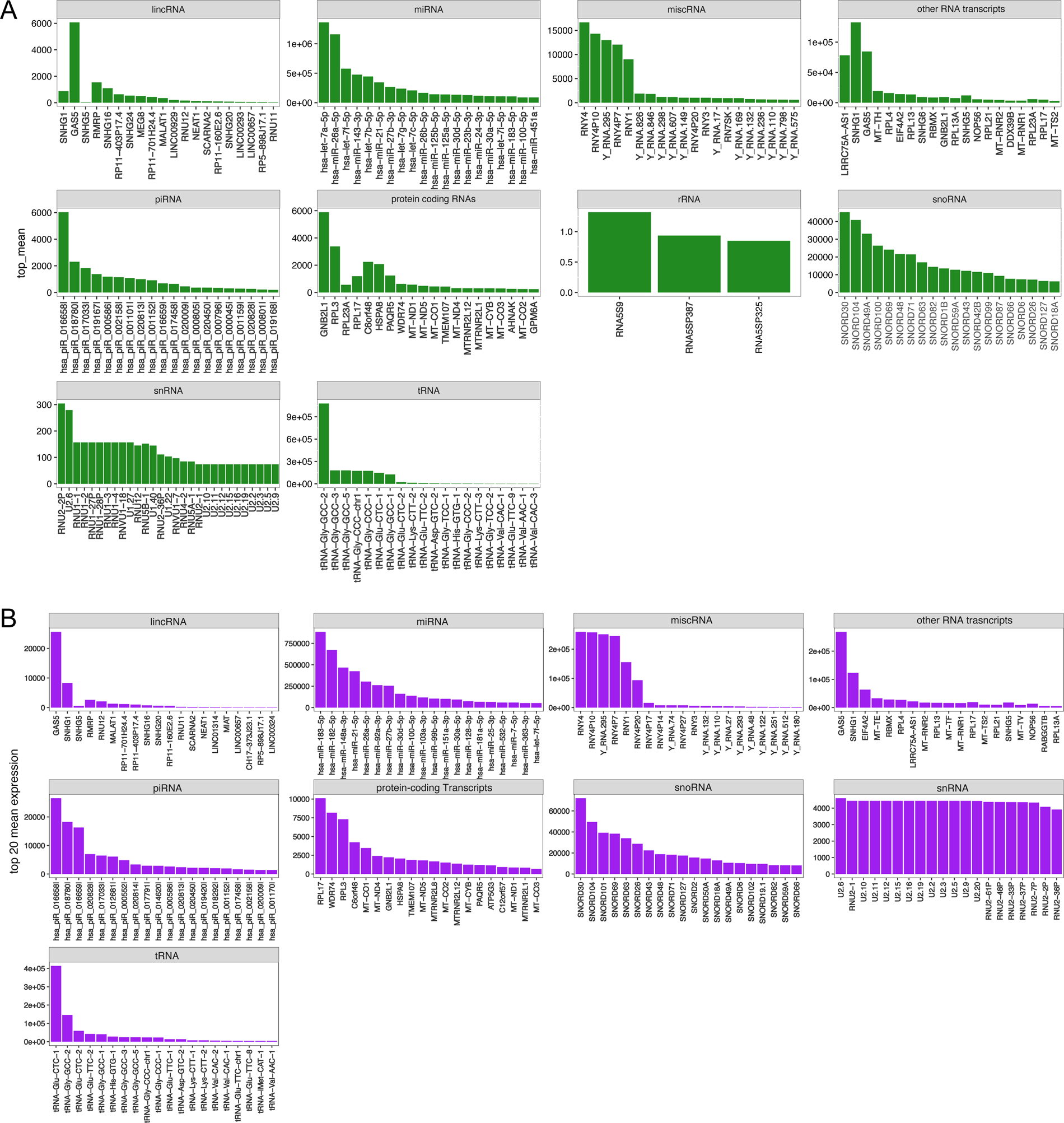
(A) Top-expressed ncRNA per biotype in hDRGs and (B) top-expressed ncRNAs per biotype in iDNs

**Supplementary Figure S4.**
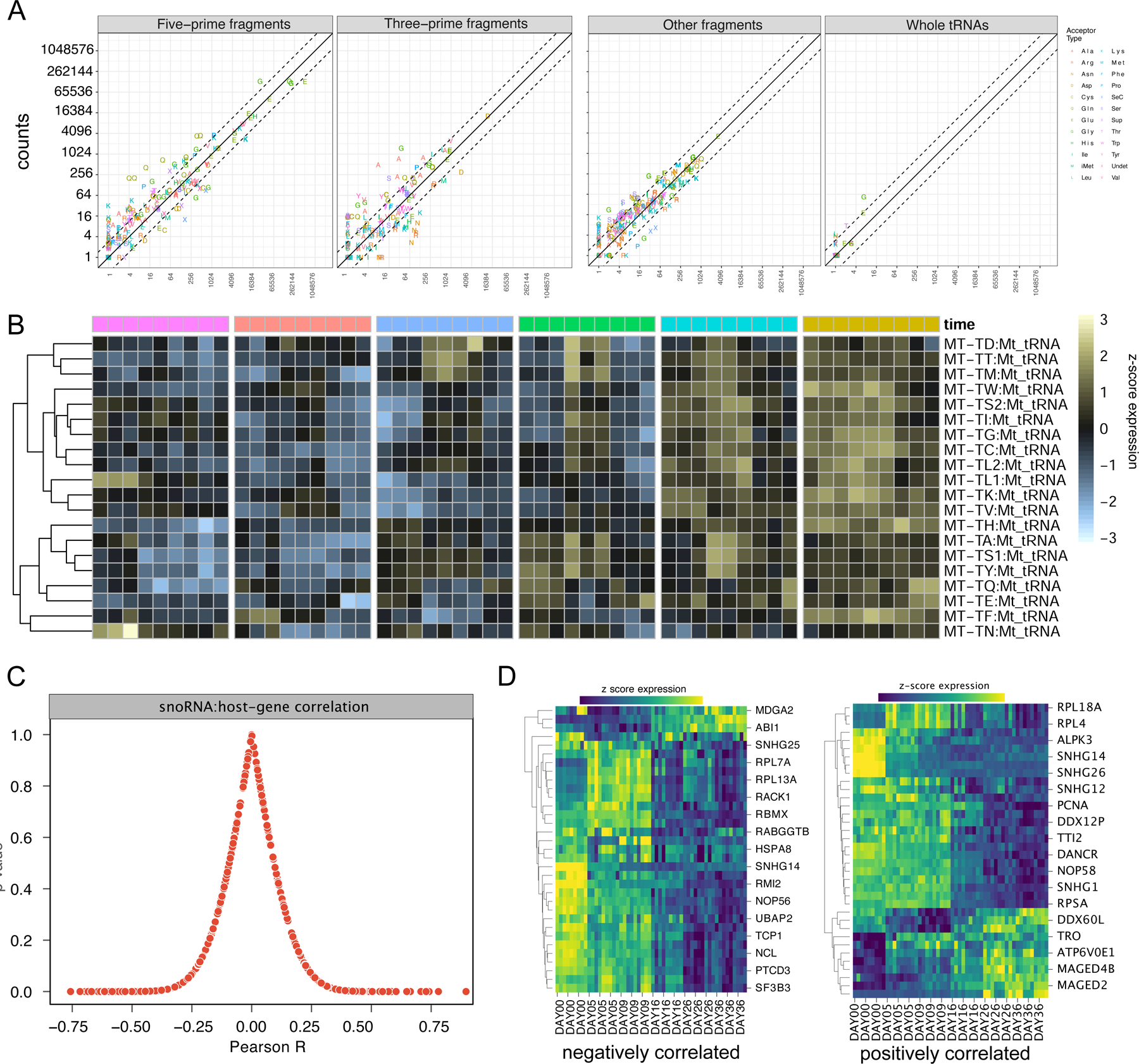
Analysis of tRNAs, mt-tRNAs and snoRNA host-genes. **(A)** tRAX was used to realign ncRNA counts to tRNAs and tRNA fragments only, since this pipeline is specialized for tRNA expression analysis. Precise location of tRNA reads were determined considering 5’ and 3’ or other fragments based on the location of the read**. (B)** Mitochondrial tRNA variance stabilized counts visualized as heatmap. **(C)** Correlation analysis of snoRNA host-genes with snoRNAs using Pearson correlation coefficient. **(D)** Expression patterns of the topmost negatively and positively correlated host-genes.

**Supplementary Figure S5.**
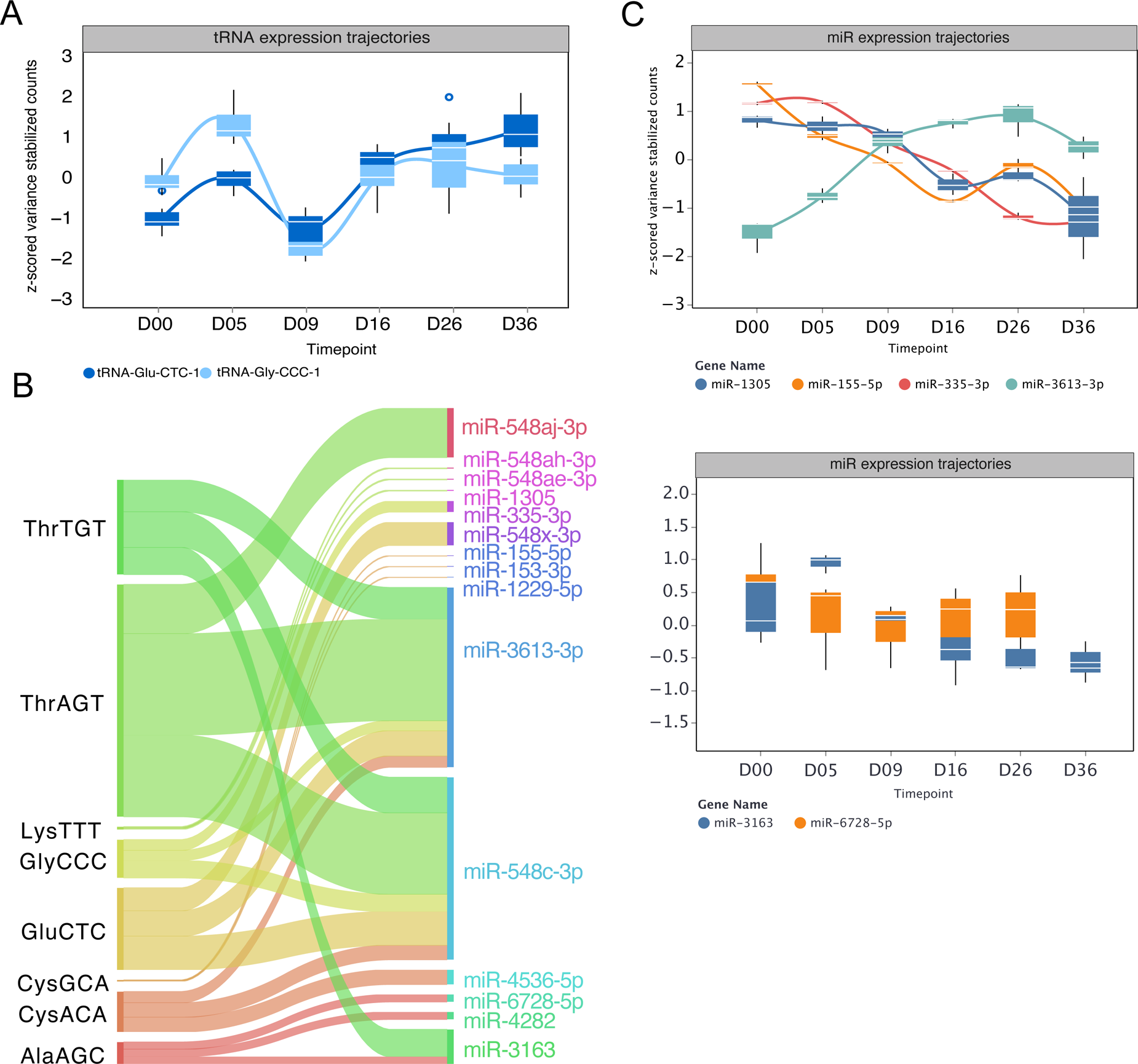
(A) Expression trajectories of tRNA-GlyCCC and tRNA-GluCTC using variance stabilized counts. Sankey plot of overlapping target-spaces between miRNAs and tRNAs indicating which miRNAs are most likely mimicked by the tRNA **(B)**. **(C)** Trajectories of mimicked miRNAs derived from NOCICEPTRA using variance stabilized counts

**Supplementary Figure S6.**
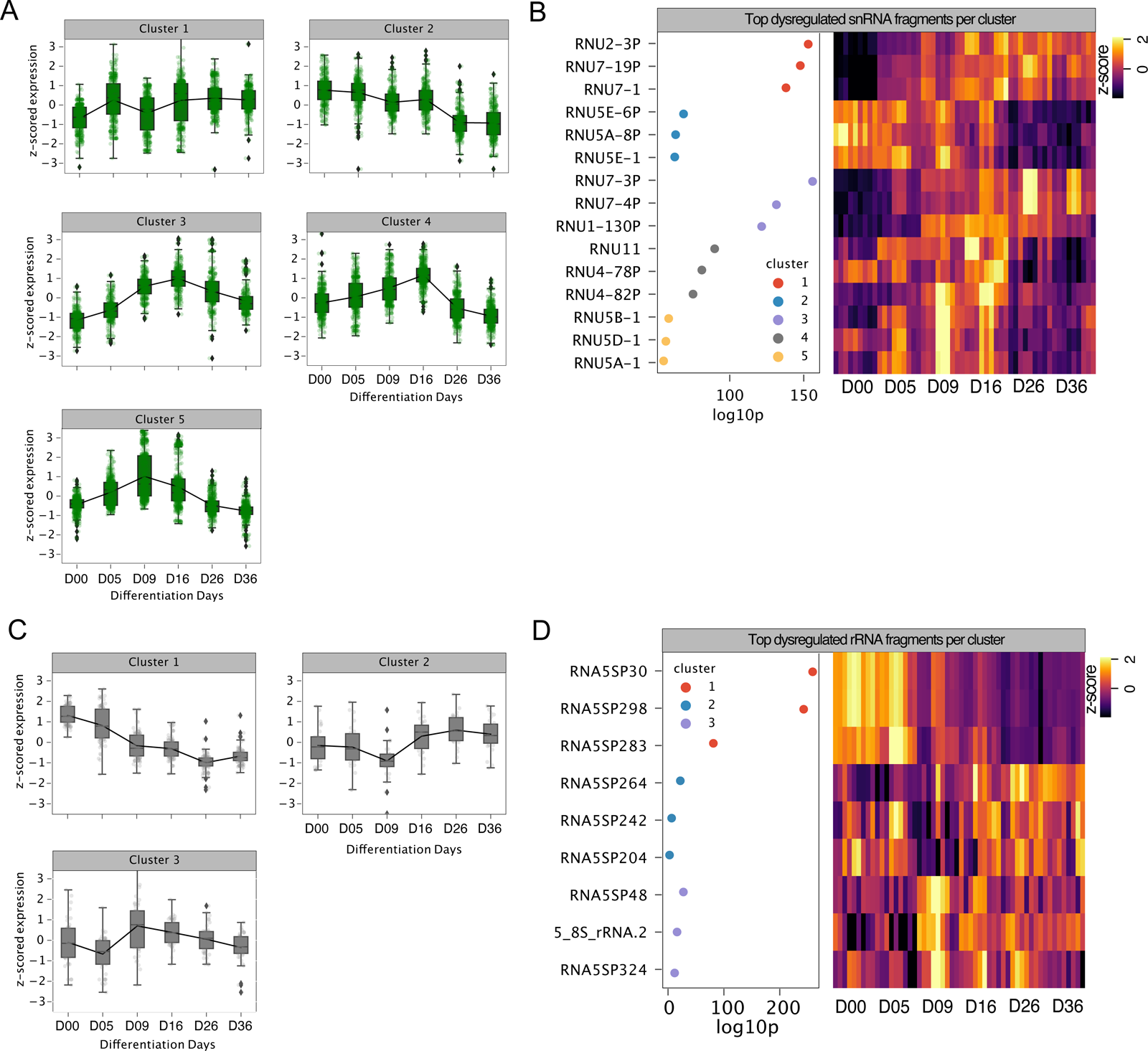
Clustering Analysis of sRNAs (A-B) and rRNAs (C-D)

## Tables

**Supplementary Table S1** Organ donor Table of hDRG samples (levels L1, L2, L3, L4, L5, T12) used to obtain Total RNAseq.

**Supplementary Table S2** Organ donor Table for hDRG samples used to obtain miRNA with the NextFlex Kit.

**Supplementary Tables 3-4** Enrichment Analysis of positively (Sheet 1) and negatively (Sheet 2) correlated snoRNA host-gene determined using g:Profiler ontology analysis.

**Supplementary Tables 5** Enrichment Analysis of tRNA target spaces.

## Notes

https://github.com/mouzkolit/Nociceptra2_Analysis

https://nociceptra.streamlit.app

### Competing Interest Statement

The authors have declared no competing interest.

### Summary of Updates

Changed the Github Links and the Weblink to the nociceptra app

